# Rethinking sinking: Imaging flow fields of natural marine aggregates to infer porosity-dependent changes in sinking velocity and carbon flux

**DOI:** 10.1101/2023.09.05.555549

**Authors:** Clara M. Flintrop, Soeren Ahmerkamp, Nasrollah Moradi, Isabell Klawonn, Jöran März, Cora Hörstmann, Rainer Kiko, Arzhang Khalili, Hans-Peter Grossart, Xosé A. Álvarez-Salgado, Javier Arístegui, Morten H. Iversen

## Abstract

The marine biological carbon pump is mainly driven by the interplay between aggregate sinking velocity and remineralization. Sinking velocity of natural marine aggregates is not routinely measured but often calculated using Stokes’ law, which does not consider size-dependent changes in porosity. We analyzed the flow fields around 81 *in situ*-formed aggregates using Particle Image Velocimetry (PIV) to determine the factors controlling aggregate settling. Using an independently derived scaling of porosity with size, we predicted the sinking velocity of laboratory-formed and *in situ*-formed aggregates with known densities. Small aggregates (<500 µm) have relatively lower porosities than large aggregates, and their increased compactness and density leads to higher size-specific settling velocities, and generally higher carbon-to-volume ratios. Applying our scaling approach to a global data set of vertical aggregate abundance and size distribution, we found that small aggregates contribute 40-70% to total carbon fluxes *in situ*.

**Teaser:** Improved sinking velocity prediction for marine aggregates highlights the contribution of small aggregates to carbon sequestration.

## Introduction

Sinking of phytoplankton cells to the deep ocean in the form of aggregated organic matter (either as phytoplankton aggregates, zooplankton fecal pellets or feeding webs, or as marine snow when exceeding 500 µm in diameter) causes an average carbon flux of 5-12 PgC yr^-1^ out of the upper 100 m of the ocean ((*1*) and references therein). The downward transport of particulate organic matter is known as the biological carbon pump (BCP; (*2, 3*), and its efficiency is strongly affected by heterotrophic organisms in the water column (*4–8*). The amount of biotic remineralization, and thus carbon flux attenuation, is strongly related to aggregate sinking velocity because it determines the residence time of the organic matter in the upper 1000 m and thereby the time available for remineralization (*7, 9–11*). Natural marine snow has high remineralization rates, and typically 80% of the particulate organic carbon (POC) contained within a slow-sinking aggregate is remineralized in the upper 200 m of the water column (*4, 12*). Despite the significance of sinking velocity for export flux, this parameter is not routinely measured. Direct measurements of aggregate sinking velocities are time-consuming and require collection of intact *in situ*-formed aggregates (*13*) or specially designed *in situ* camera system (*14*). Several methods are used to measure aggregate sinking velocity, e.g., net-jet flow systems (*9, 15, 16*), sinking columns (*17–19*), and roller tanks (*20, 21*). Because of restrictions imposed by sampling depth and processing time, *in situ* camera systems like the Underwater Velocity Profiler (UVP; (*6, 22*)) have become the gold standard for high-throughput vertical profiling of aggregate abundance -, volume -, and size-spectra (see (*23*) for a comprehensive overview of optical systems). Although optical systems do not routinely measure sinking velocity, they are often used for carbon flux calculations by assuming that aggregate size scales as a power function to both sinking velocity and mass (*7, 24, 25*).

To indirectly determine carbon flux in the water column using *in situ* camera profiles, aggregate sinking velocity is often calculated according to Stokes’ law (*26, 27*), where sinking velocity is a function of the squared aggregate radius and the excess density. However, it has been observed that calculations based on Stokes’ law tend to underestimate the sinking velocity of smaller aggregates while overestimating the velocity of larger aggregates (*16, 28*). Several properties of marine snow, including its highly variable morphology, surface roughness, and porosity, have been considered to explain the non-compliance with Stokes’ law (*29–35*). Deviation from Stokes’ law due to aggregate heterogeneity makes it difficult to determine a global relationship between size and sinking (e.g., (*7, 36*)). Still, if aggregate density and porosity is considered, we hypothesize that general relationships can be identified when testing Stokes’ sinking for different types of aggregates. We used Particle Image Velocimetry (PIV) to visualize the flow field around *in situ*-formed aggregates and laboratory-made impermeable agar spheres, which we used as reference for Stokes-compliant aggregates. To determine the factors controlling aggregate sinking velocity and, thus, improve existing parameterizations, we compared measured sinking velocities to Stokes’ prediction, and related sinking behavior to the hydrodynamic flow field and physical parameters (size, porosity, density, structure, and composition) of agar spheres and *in situ*-formed aggregates. We derived an independent scaling of porosity with size, which allowed accurate prediction of size-specific sinking velocities of both laboratory-made and *in situ*-formed aggregates with known densities. This improved carbon flux predictions considerably compared to those derived via Stokes’ law, which underestimated carbon flux by 2-fold, on average. Our results allow parameterization of organic aggregates in Earth system models and improve carbon flux estimates to better predict future changes in the biological carbon pump following climate change-induced shifts in phytoplankton community structure.

## Results

### Aggregate type, size and sinking velocity

The study was carried out in July and August 2017 in the North-West African Eastern Boundary Upwelling System during the campaign “Fluxes I” on board B/O Sarmiento de Gamboa (Fig. 1, A). During the expedition, we collected a total of 108 *in situ*-formed aggregates using a Marine Snow Catcher (see “Aggregate collection” in the Materials and Methods section), which ranged in size from 0.38 mm to 3.56 mm (mean 1.15 (SD 0.65) mm) in equivalent spherical diameter (ESD; Table 1). The PIV measurements for 27 aggregates did not pass quality control (see ‘Particle Image Velocimetry’ in the Materials and Methods section), leaving 81 aggregates for further analysis. Based on their main constituents, we assigned them to three separate categories: phytoplankton aggregates, degraded fecal pellets, and ballasted mucus aggregates (Fig. 1, B). Phytoplankton aggregates made up 45% of the total aggregate numbers and were distinguished from other aggregates by their heterogeneous composition, which included diatom chains, Acantharia, Radiolaria, mucilaginous aggregates from discarded zooplankton feeding webs, fresh and degraded fecal pellets, copepod carcasses, and other unidentified phyto- and zoo-detritus. Another 28% of the total aggregates were categorized as fecal pellets at various stages of remineralization based on particle size, homogeneous distribution of matter, uniform coloring, and their oval shape (from here on also referred to as “fecal pellets”). A further 27% were categorized as ballasted mucus aggregates as they were comprised of a translucent, mucilaginous substance, possibly originating from discarded zooplankton feeding webs. Average gray value distributions of grayscale images of aggregates taken under the microscope showed distinct differences between the three categories, allowing a semi-quantitative classification of aggregates from these distributions alone (Fig. S1, A). Sinking velocities, which were measured for individual aggregates in a net-jet flow system (see “Aggregate properties” in the Materials and Methods section) ranged from 50 to 410 m d^-1^ (mean 156 m d^-1^ ±74 SD; Table 1), which is comparable to other measurements in the region (Fig. S2, A; Data S1; (*37–39*)). Agar sphere sinking velocity increased linearly with size, but there was no apparent correlation between size and sinking velocity for field*-* collected aggregates, even after categorizing aggregates based on their composition (Fig. S2, B). In addition to measuring sinking velocity experimentally, we also calculated theoretical sinking velocities according to basic force balance equations (Eqs. M1-M6; all equations found in the ‘Materials and Methods’ section have the letter ‘M’ preceding the equation number) and found that calculated sinking velocities deviated from measured sinking velocities by an average of 30%, even when adjusting the drag coefficient according to (*40*) (Eq. M7).

**Table 1.** Overview of measured parameters for the different aggregate types (mean±standard deviation): Number of aggregates *n* per category and parameter (except for *ρ*_*shd*_), aggregate diameter *d*, sinking velocity *w*, Reynolds number *Re*, drag coefficient *C*_*D*_, excess density *Δρ*, solid hydrated density *ρ*_*shd*_ (number of aggregates for which parameter was measured in parentheses), and porosity *φ*. ‘All *in situ*-formed aggregates’ denote the sum of *in situ*-formed aggregates used for further analysis, excluding agar spheres.

| Parameter | All *in situ*-formed aggregates | Phytoplankton aggregates | Ballasted mucus aggregates | Degraded fecal pellets | Agar spheres |
| --- | --- | --- | --- | --- | --- |
| $n$ | 81 | 36 | 22 | 23 | 13 |
| $d$ (mm) | 1.17±0.61 | 1.36±0.77 | 1.21±0.44 | 0.84±0.20 | 1.26±0.31 |
| $w$ (m d<sup>-1</sup>) | 146.1±56.3 | 147.2±68.0 | 163.4±48.3 | 127.8±35.9 | 146.2±60.6 |
| $Re$ | 1.8±1.1 | 2.1±1.3 | 2.1±1.1 | 1.2±0.4 | 2.1±1.3 |
| $C_D$ | 22.0±12.2 | 21.6±14.4 | 17.1±6.6 | 27.6±10.5 | 20.0±12.3 |
| $\Delta\rho$ (g cm<sup>-3</sup>) | 0.0051±0.0046 | 0.0053±0.0061 | 0.0044±0.0027 | 0.0056±0.0030 | 0.0027±0.0004 |
| $\rho_{shd}$ (g cm<sup>-3</sup>) | 1.24±0.09 (40) | 1.17±0.04 (17) | 1.26±0.05 (9) | 1.31±0.08 (14) | 1.14±0.02 (7) |
| $\varphi$ | 0.973±0.03 | 0.963±0.04 | 0.981±0.01 | 0.981±0.01 | 0.976±0.003 |

**Fig. 1.**
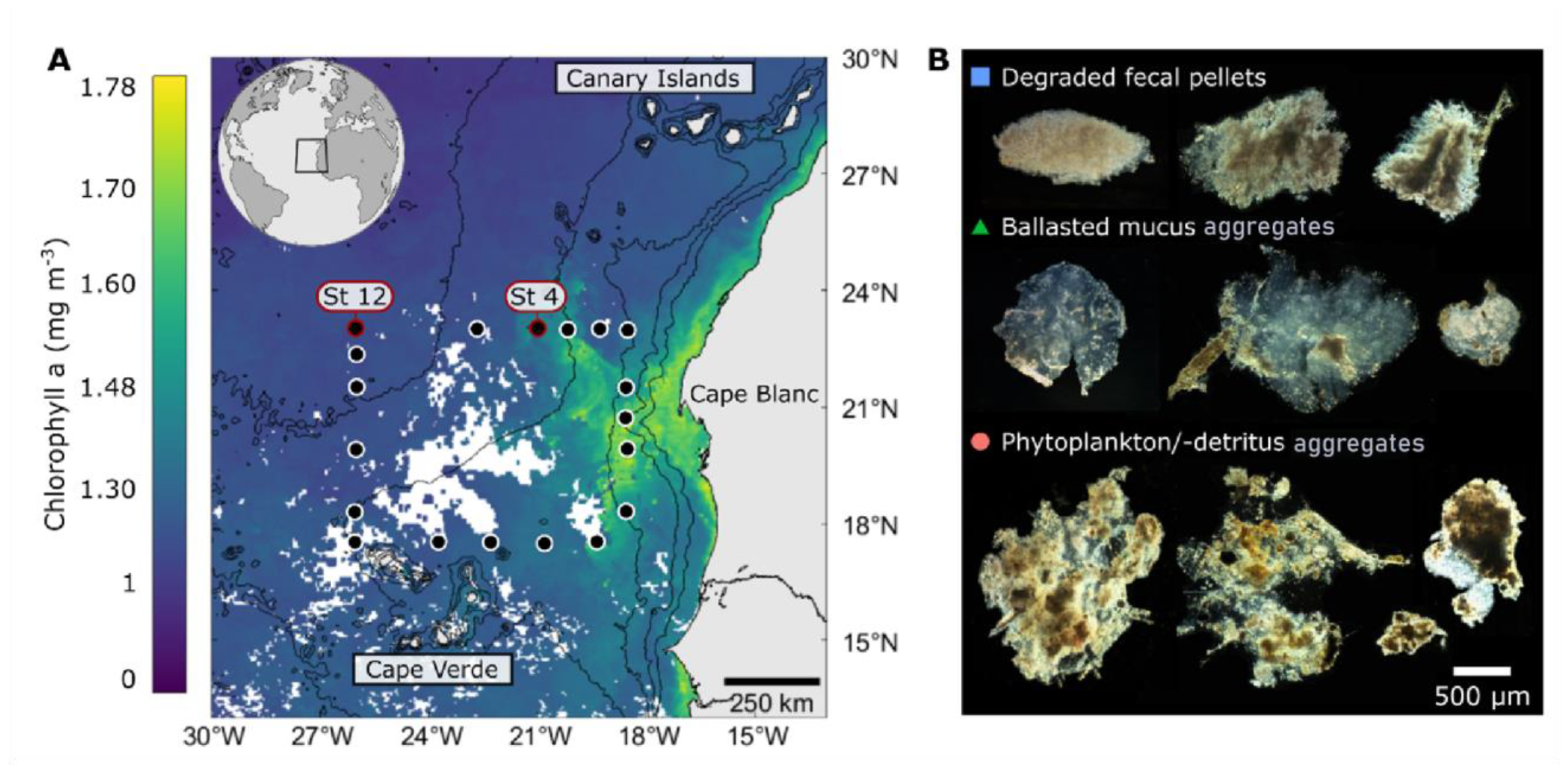
Study area. (**A)** Sampling stations (black dots) and sea surface chlorophyll a concentration during the sampling period. Stations where sediment traps were deployed are circled in red. (**B**) Examples of the different types of aggregates formed *in situ*.

### Flow fields around sinking marine snow

We analyzed the flow fields around 81 *in situ*-formed aggregates of varying composition and structure using Particle Image Velocimetry (PIV; Movies S1) (the PIV recordings of 12 out of the 93 categorized aggregates did not pass quality control, see “Particle Image Velocimetry” in the Materials and Methods section). As reference for Stokes-compliant aggregates, we additionally analyzed the flow fields around 13 agar spheres representing perfectly spherical, impermeable spheres. To compare the flow fields around aggregates with different sinking velocities and sizes, we calculated their Reynolds numbers (*Re*), which ranged from 0.4 to 5.2 (mean 1.9 (SD 1.2); Table 1). This Reynolds number regime represents an intermediate range above creeping flow conditions (*Re* << 1) and far below a turbulent regime (*Re* > 2300). For *Re* ∼ 1, the flow field was mostly symmetric, i.e., the velocity boundary layer thickness at the upstream side of the aggregate equaled that at the downstream side, with only small variations. With increasing *Re*, the flow field became increasingly asymmetric. Overall, at similar *Re*, the flow fields of agar spheres and marine aggregates of various shapes and compositions were quite similar (Fig. 2, A-C).

**Fig. 2.**
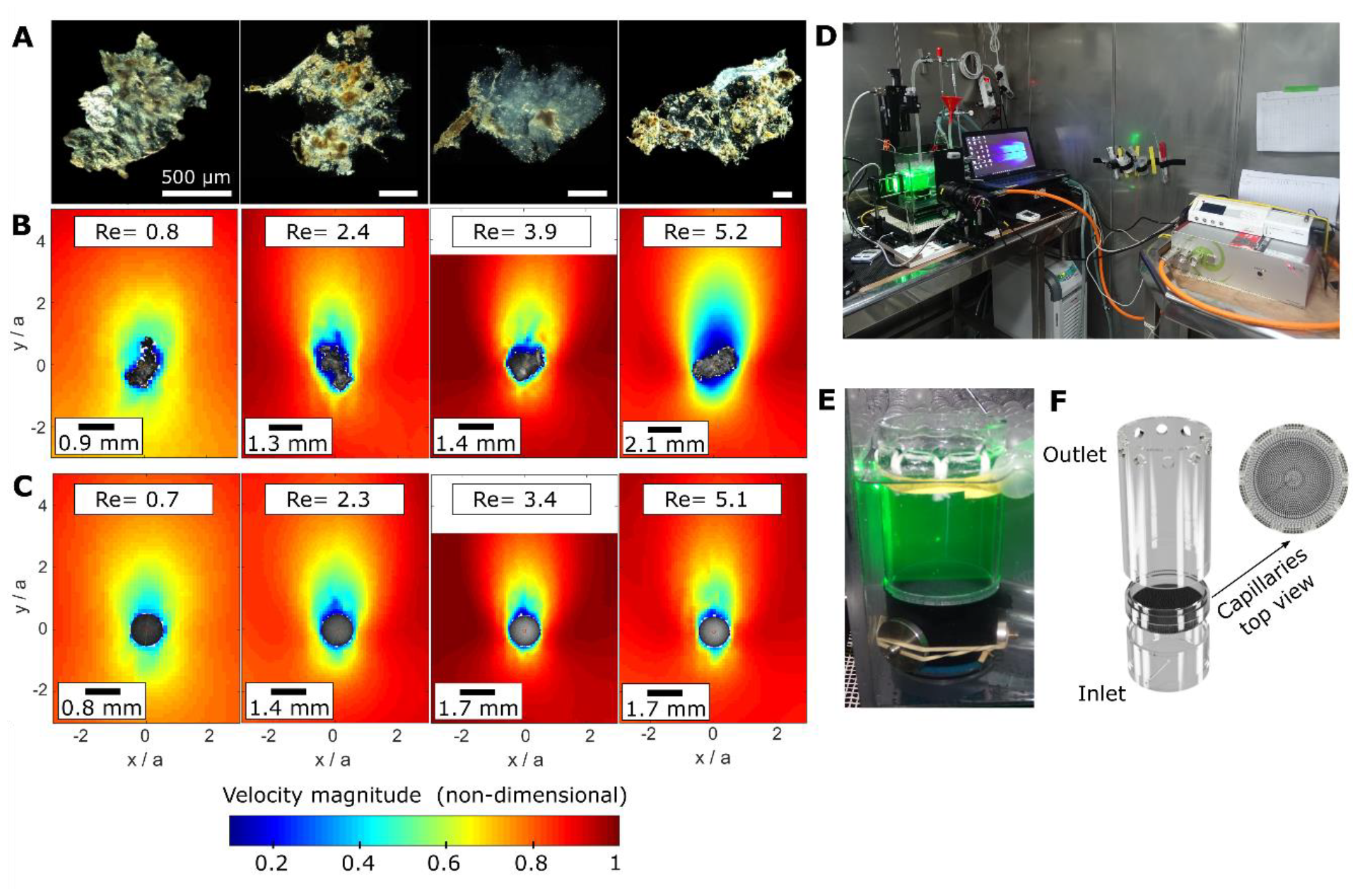
Aggregate flow fields and PIV set-up. (**A**) *In situ*-formed aggregates with their corresponding flow fields (**B**). (**C**) Flow field comparison with agar spheres with comparable Reynolds number. Note that the flow fields around the last two agar spheres look very similar despite their Reynolds numbers differing by a factor of 1.5, due to slightly different excess densities of the individual agar spheres. (**D**) Flow chamber set-up on board B/O Sarmiento de Gamboa. (**E**) Aggregate is freely suspended in the flow chamber and illuminated by a pulsed light sheet (green) for PIV. (**F**) Modified design of the flow chamber for optimized laminar flow.

To quantify variations in the flow field for different *Re* regimes, we measured the thickness of the velocity boundary layer at the aggregate equator, defined as the fluid layer at the aggregate surface where flow velocity was ≤ 90% of the bulk flow velocity. Scaled boundary layer thickness, i.e., boundary layer thickness normalized by the x-axis diameter (*δ*_*s*_ = *δ*/*d*) ranged from 0.37 to 2.11 (Fig. 3, A), and increased the apparent aggregate diameter up to 5-fold. There were no obvious patterns of deviation from the regression lines fitted for agar spheres and the *in situ*-formed aggregates, except that with increasing *Re* (starting around *Re* = 3), the boundary layer of aggregates became relatively thinner compared to that of the agar spheres (Fig. 3, A). Irrespective of aggregate type, the scaled boundary layer thickness scaled with *Re* following a power law (*δ*_*s*_ = 1.37·*Re*^-0.46^). *In situ*-formed aggregates had irregular shapes and surfaces (Fig. 1, B; Fig. S3), but there was no obvious difference between the power law coefficients for agar spheres (*δ*_*s*_ = 1.45·*Re*^-0.48^) and any of the aggregate types (Fig. 3, A). To analyze the extent to which the deviation between theory and our measurements affected sinking velocity prediction, the drag coefficient *C*_*D*_ was estimated based on the measured boundary layer thickness (Eq. M9), and plotted against *Re*, giving Eq. 1. Generally, the drag coefficients for marine aggregates and agar spheres scaled with *Re* as expected from Stokes’ law (Fig. 3, B) with a best fit of

**Fig. 3.**
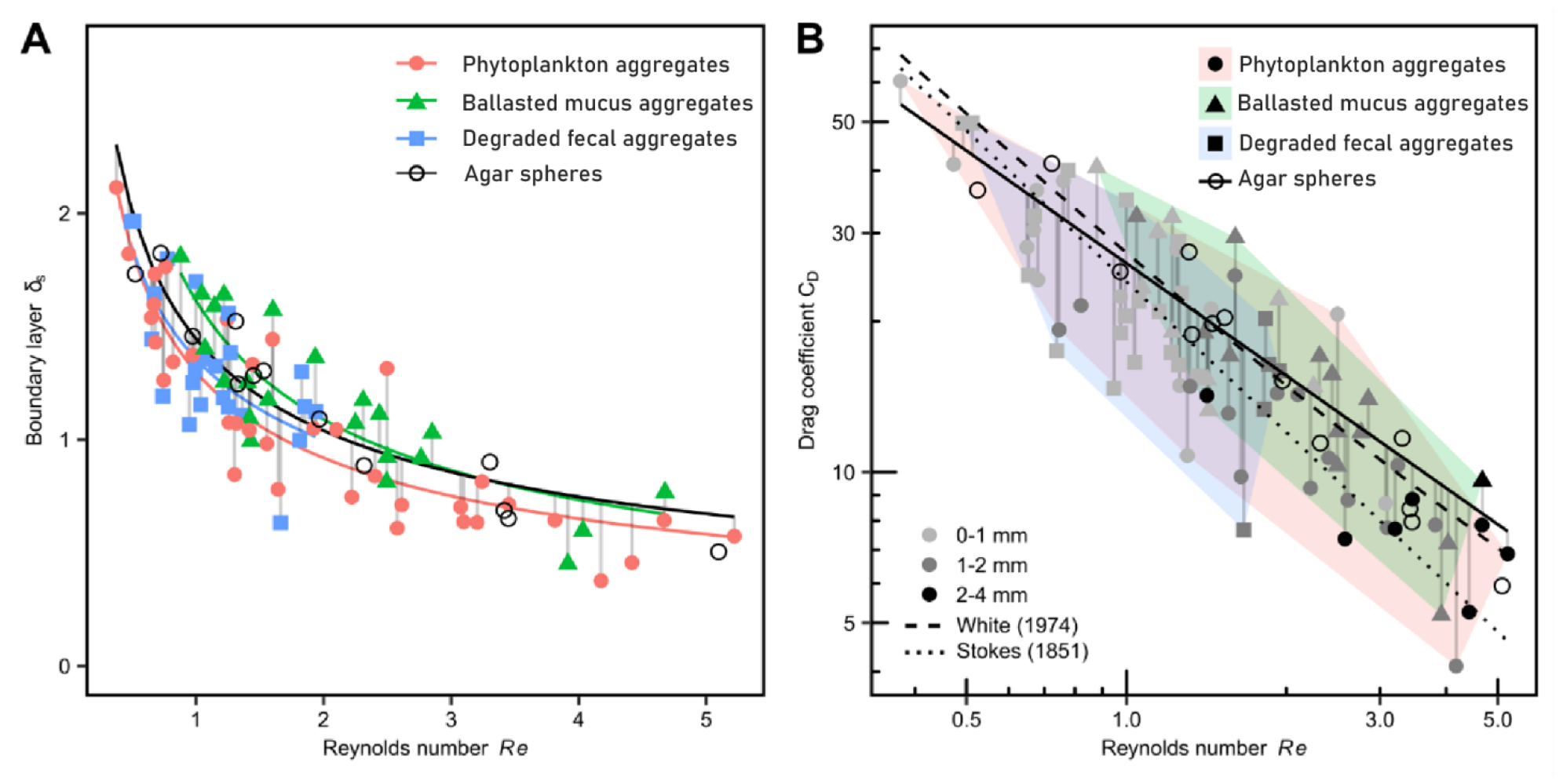
Reynolds number scaling. Relationship between Reynolds number and (**A**) scaled equatorial boundary layer *δ*_*s*_, and (**B**) *δ*_*s*_–based drag coefficient *C*_*D*_ (Eq. M9) for the different aggregate types and agar spheres, plotted on a log-log scale. The solid black lines in panels **A** and **B** denote regression lines fitted to the agar spheres, which represent Stokes-compliant (i.e., perfectly round and impermeable) particles. In panel **B**, the regression lines obtained using the Stokes equation ((*27*); Eq. M6, dotted line) and the Stokes equation with a modified drag coefficient according to White ((*40*); Eq. M7, dashed line) are also shown for comparison.

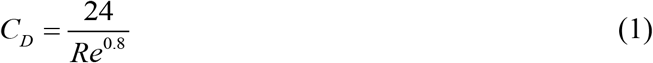

Residuals between estimated drag coefficients for *in situ*-formed aggregates and drag coefficients predicted according to Stokes’ law (using agar spheres as reference) were calculated as (*C*_*D Aggregates*_ – *C*_*D Agar spheres*_) / *C*_*D Agar spheres*_ using Eq. M9. Residuals were largest for small Reynolds numbers (*Re* < 1.5), i.e., large boundary layer thickness, where the drag coefficient is underestimated by Stokes’ law (up to 5-fold, on average almost 2-fold). This deviation translates into an overestimation of the sinking velocity by 40-120% when calculated using Stokes’ law. For larger Reynolds numbers (*Re* > 1.5), the residuals strongly decreased in magnitude (up to 2-fold, on average 1.5-fold), translating into sinking velocity overestimation of 20-40% when calculated with Stokes’ law. By following the White extension for the drag coefficient ((*40*); Eq. M7), the residuals further decreased by an average of 30% across the whole Reynolds number range and the trend towards overestimation of sinking velocity for large aggregates was diminished. The magnitude of the residuals could not be explained by any of the measured individual aggregate properties. Residuals varied between the different aggregate types, showing that Stokes’ Law was better at predicting the sinking velocity of some aggregate types, especially at low Reynolds numbers (*Re* < 1.5) range. Of all the shape factors considered (see “Aggregate properties” in the Materials and Methods section), only aspect ratio and orientation had some influence on the magnitude of the residuals, but for the majority of shape factors, the effect was not statistically significant due to the large scatter amongst aggregates of similar types (Fig. S3; significance tested using a t-test, with p-values given for slopes that differ statistically significantly from zero). The average error in predicting sinking velocity based on the White drag coefficient was estimated to be 10% for degraded fecal pellets, 16% for phytoplankton aggregate, and 26% for ballasted mucus aggregates.

### Excess density, porosity, and fractality

The excess density of aggregates depends on the density of the solid fraction (solid hydrated density *ρ*_*shd*_) and the porosity *φ* relative to the water density *ρ*_*w*_. While solid hydrated density could be measured directly, we estimated the excess density using Eq. M8, where the boundary layer-estimated drag coefficient (Eq. M9) was used. We found the excess density to range over two orders of magnitude from 0.0002 g cm^-3^ to 0.0209 g cm^-3^ (Table 1). Measured solid hydrated densities ranged from 1.119 g cm^-3^ to 1.430 g cm^-3^, spanning a considerable range of solid hydrated density measurements from the literature for organisms and aggregates of various origins (Table S1). Yet, they only explained a minor fraction of the variability in excess densities (Table 1). Porosity (Eq. M10) ranged from 0.859 to 0.999, pointing to a large water fraction in the interstitials of the aggregates (Table 1). The volume fraction made up by solid material ranged from 0.141 to 0.001, encompassing two orders of magnitude and thereby explaining most of the observed variability of the excess densities. The amount of the solid fraction was related to the diameter of the aggregate by a power law function (Fig. 4, A), giving Eqs. 2.a-2.c depending on the aggregate type considered:

**Fig. 4.**
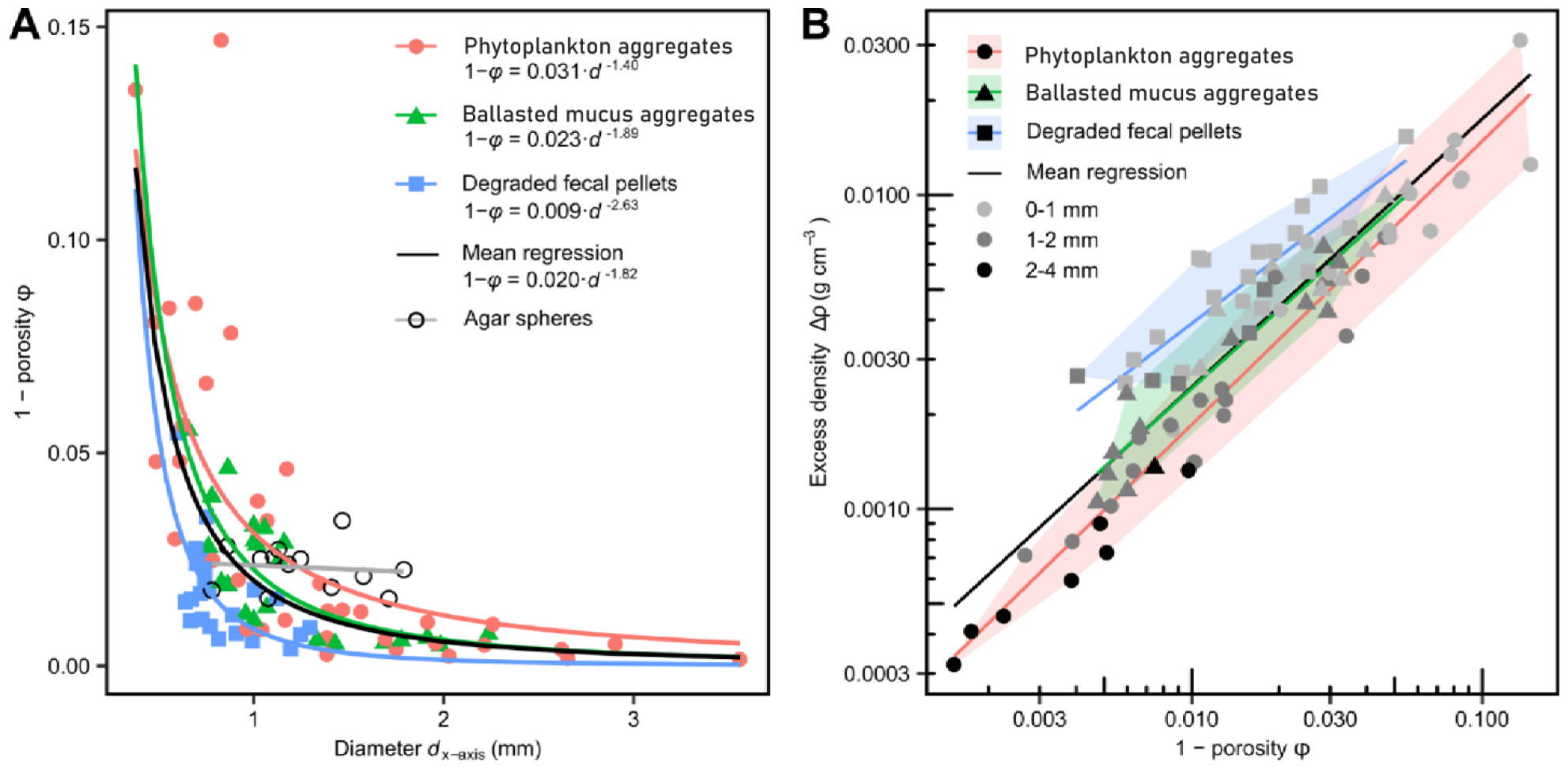
Porosity scaling. (**A**) Relationship between aggregate solid fraction and aggregate size for different aggregate types and agar spheres. (**B**) Relationship between aggregate solid fraction, aggregate excess density and aggregate size for different aggregate types, plotted on a log-log scale.

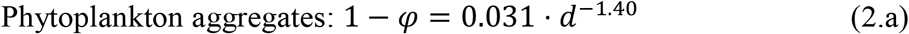

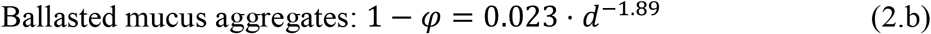

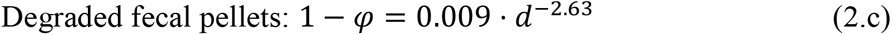

This implies that larger aggregates had an increased porosity, and therefore a reduced excess density (Fig. 4, B). This behavior is expected when taking fractal theories into account. The 1-dimensional fractal number (DF1) (inferred from the log-log relationship between perimeter and maximum aggregate length (*41*)) indicates the roughness of the outer structure relative to a smooth sphere. Values around unity indicate a smooth surface, while larger numbers indicate uneven surfaces with structures that may reach into the boundary layer surrounding the aggregates. Both agar spheres and ballasted mucus aggregates showed smooth outer structures with DF1 values of 1.18 and 1.11, respectively. Degraded fecal pellets and phytoplankton aggregates had a higher surface roughness with a DF1 value of 1.47. The 2-dimensional fractal number (DF2) (inferred from the log-log relationship between maximum aggregate length and area (*41*)) relates the area of the aggregate to the area of a sphere with the same diameter. Values of DF2 varied between 1.62 and 2.00, with ballasted mucous aggregates having the lowest values, possibly due to their relatively higher plasticity leading to deformation during sinking (Table S2). The excess density (calculated using Eqs. M5, M7, and M8) could be best fitted to the porosity (calculated using Eqs. M8-M10), as described by Eq. 3:

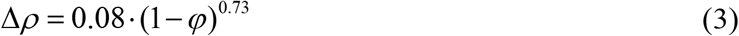

### *In situ* aggregate carbon content and fluxes

To obtain the *in situ* carbon content of marine snow, we deployed drifting sediment traps at water depths of 100 m and 200 m for 22 hours (Table S3). The first drifting traps comprised two gel traps per depth to non-destructively collect aggregates, enabling us to determine the carbon fluxes and size-spectrum of the field*-*collected marine snow (Fig. S4). The *in situ* total organic carbon fluxes were relatively constant at 42 mg C m^-2^ d^-1^. To determine the size-specific carbon content of marine snow, we utilized the gel-trap aggregate numbers from specific size bins (see Material and Methods) and calculated the Volume *V* (mm^3^) as a function of equivalent circular diameter *d*, 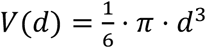, assuming a spherical shape for the aggregates. We then applied our size-dependent porosity (*φ*) relationship (Eq. 2) to estimate the solid fraction of the biovolume *B*. By assuming that all the measured carbon is equally distributed, we determined the relationship between carbon content and biomass (Eq. 4):

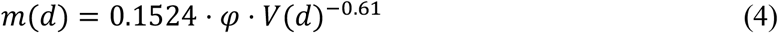

where *m*(*d*) is carbon content (µg mm^-3^). When expressed as aggregate-specific carbon-content (µg C aggregate^-1^) per size bin, this relationship changes to *m*(*d*)*∝V*(*d*)^0.39^which is slightly lower than a previously determined scaling of *m*(*d*)*∝V*(*d*)^0.52^ based on direct carbon measurements of single aggregates from the Peruvian upwelling region (*42*).

### Regional and global sinking velocity and carbon flux predictions

The newly derived size-dependent relationships for sinking velocity (Eq. M11) and carbon content (Eq. 4) provide a means to determine i) porosity *φ* and—if solid hydrated density *ρ*_*shd*_ is measured or known from literature (Table S1) —sinking velocity *w* based on aggregate size, and ii) carbon fluxes based on *in situ* measured size spectra. To test the validity of our model, we compared measured and predicted sinking velocities of aggregates with known solid hydrated density formed in roller tanks (*43–46*) and collected in the field at different locations (*47*), including aggregates collected on our research expedition (Fig. 5, A; Table S4). We observed large variabilities in predicted sinking velocities (*w*_*predicted*_) versus measured sinking velocities (*w*_*measured*_), with the predicted sinking velocity being overestimated up to 3-fold for some aggregates, and underestimated up to 5-fold for others. Still, we found a similar or better correlation between measured and predicted sinking velocities of aggregates across all datasets combined (*w*_*predicted*_ = 0.78·*w*_*measured*_ +104, adj. R2 = 0.33, or *w*_*predicted*_ = 1.19·*w*_*measured*_, adj. R2 = 0.17, when the intercept is forced through zero) compared to previously established relationships, e.g., as compiled in (*48*) (Fig. S5).

**Fig. 5.**
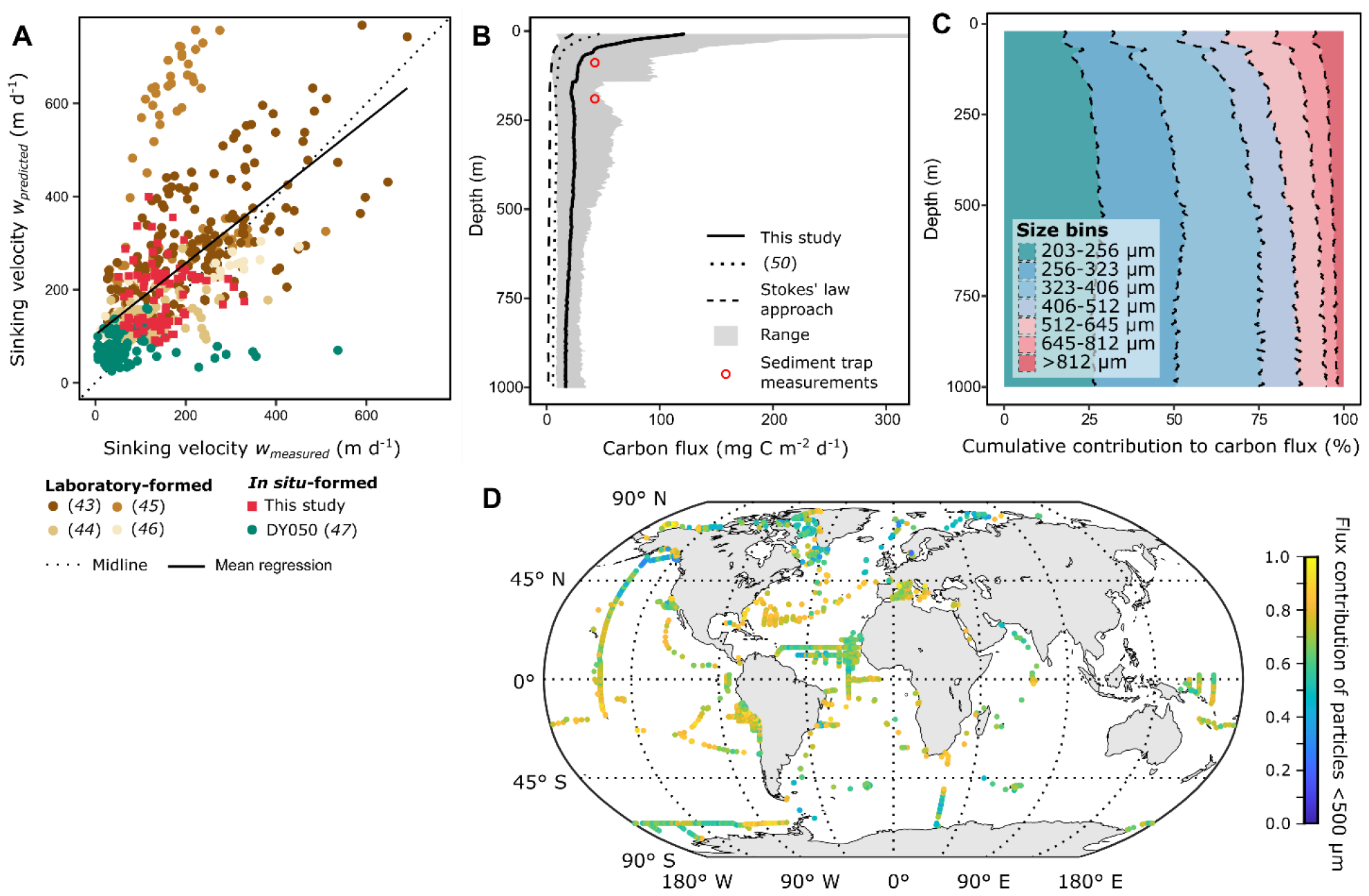
Sinking velocity and carbon flux predictions. (**A**) Measured versus predicted sinking velocity of aggregates from this study, as well as selected laboratory and field datasets (Table S4). Predicted sinking velocities were calculated using *w* = *d*^2^ *⋅ g ⋅* (*k ⋅ d*^*p*^ *⋅ 𝜌*_*shd*_ + *𝜌*_*w*_ *⋅* (*k ⋅ d*^*p*^ ™ 2))*/(18η*) (Eq. M11), where *k* is the coefficient and *p* is the exponent of Eq. 2.a. (**B**) Carbon flux calculated using our relationship (solid line), the relationship used in (*50*) (dotted line) and Stokes’ law approach (dashed line) compared to measured sediment trap fluxes at 100 m and 200 m. (**C**) Cumulative contribution of aggregate size classes to total carbon flux. (**D**) Relative contribution of small aggregates (<500 µm) to total carbon flux at the base of the mixed layer.

To test the validity of our carbon content and carbon flux predictions, we averaged 72 vertical profiles of *in situ* aggregate abundance and size-distribution made with a UVP system during the same cruise and applied our relationships (Eqs. 2, 4, and M13) to calculate carbon fluxes and assess the contribution of specific size classes to the overall carbon flux. The overall carbon flux at 100 m and 200 m was 24 mg C m^-2^ d^-1^ and 35 mg C m^-2^ d^-1^, respectively, which is on average 2-fold higher than predicted based on Stokes’ law, but consistent with the sediment trap data (Fig. 5, B). Intriguingly, our estimations indicate that small aggregates (<500 µm) contribute a remarkable 80% of the overall carbon flux (Fig. 5, C). This proportion is substantially higher than the 9% predicted by Stokes’ law, highlighting the significant role of abundant small aggregates for ocean carbon fluxes. To assess the impact of small aggregates on carbon fluxes on a global scale, we conducted a thorough analysis utilizing 10,907 UVP-based aggregate profiles obtained from the publicly available ECOTAXA database (see ‘Material and Methods’). Carbon fluxes were determined at the annual mean mixed layer depth (*49*) using Eq. M12, ranging from 2.3 mg C m^-2^ d^-1^ (10^th^ percentile) to 12.7 mg C m^-2^ d^-1^ (90^th^ percentile) on a global scale. Our findings revealed that small aggregates, on average, contribute 67% to the global carbon flux at the base of the mixed layer (*49*) (Fig. 5, D). Notably, distinct meridional differences were observed, with the highest flux contribution of small aggregates occurring in the region between 20°N-20°S (81-85%) and a still substantially large contribution of 46-50% at mid-latitudes between 20°N-65°N and 20°S-65°S.

## Discussion

### Improved porosity measurement and sinking velocity prediction

We used four independent measurements (i) aggregate size *d*, (ii) sinking velocity *w*, and (iii) normalized velocity boundary layer thickness *δ*_*s*_ to estimate the drag coefficient *C*_*D*_, plus (iv) solid hydrated density *ρ*_*shd*_ to get an improved measurement of aggregate porosity in relation to particle size. We collected aggregates with a wide range of compositions, sizes, and sinking velocities that reflect the diversity and heterogeneity of *in situ*-formed aggregates and match previously reported values of size and sinking velocity for natural marine aggregates. Visualization of the flow field around aggregates showed that the fluid boundary layer thickness, drag coefficient, and general flow field did not noticeably differ from that of impermeable spheres with similar Reynolds numbers, suggesting that *in situ-* formed aggregates behave according to basic force balances. Deviations in measured sinking velocities seem to stem from the porosity of marine aggregates, which increases with increasing aggregate size following a power function (*8, 9, 32, 45, 51*). The increased porosity for large aggregates caused their excess density to decrease noticeably with increasing diameter. This resulted in lower sinking velocities for large aggregates than solely predicted by Stokes’ drag (*52*). The drag coefficient estimated from the boundary layer thickness scaled with the Reynolds number. As expected from the Reynolds number regime (*Re* = 0.4-5.2), the Stokes’ drag 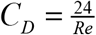 (*27*) was not sufficient to predict sinking velocity. The White relationship (Eq. M7; (*40*)) as well as our best fit of *C*_*D*_ = 24 /*Re*0.8 (Eq. 1) yielded a significant improvement. A similar fit was also found through theoretical approximation by (*53*) (*C*_*D*_ = 29.03/*Re*^0.87^). By applying the drag as well as the porosity relationships, the controlling variables can be reduced to diameter, solid hydrated density, gravitational acceleration, water viscosity, and water density. Knowing these parameters, the sinking velocity of marine aggregates can be calculated in a straightforward manner by combining Eqs. 2.a, 3, and M6 (resulting in Eq. M11) as presented in Fig. 5, A.

### Importance of aggregate composition and density

Even though the new parameterization predicts the magnitude and trends of sinking velocities well, large uncertainties were observed. These were mainly caused by large variability in the solid hydrated density (Table 1), showing that despite the comparatively low content of solids in marine aggregates, solid hydrated density contributes significantly to the sinking behavior of marine snow. Solids can affect sinking of marine aggregates in two major ways: the density of solid components affects aggregates such as marine snow by increasing their mass, i.e., excess density (*52*). At the same time, shape and properties of solids affect how tightly they can be packed within an aggregate (i.e., the compactness) and directly affect the porosity of an aggregate (*9, 54*). Average solid hydrated density of aggregates is likely determined by the make-up of the phytoplankton community (*9, 34, 55*) and lithogenic ballasting minerals (*8, 45*). We know that composition and compactness affect aggregate density, but it remains a task for future studies to further investigate the regulatory mechanisms for solid hydrated density and derive global relationships. Until then, literature values can be used to infer sinking velocity from our global porosity scaling (Table S1). Furthermore, we did not find any indication that irregularities in aggregate shape substantially influenced the drag coefficient, even when comparing between aggregate types and compared to agar spheres (Fig. S3). This supports previous studies that found shape, permeability, and surface roughness to be poor predictors of sinking velocity (*29–33*).

Overall, we did not find any difference in physical properties, including sinking behavior, between *in situ-*formed aggregates and impermeable agar spheres, except that agar spheres are not fractal and, therefore, porosity did not increase with increasing agar sphere size (Fig. 4, A). This implies that the sinking velocity of marine aggregates can be predicted via the basic force balance equation once the solid hydrated density is known and the size-porosity relationship is taken into account. Furthermore, the flow fields around field-collected aggregates and agar spheres had matching magnitudes and symmetries at similar Reynolds numbers (Fig. 2, B, C). Although we cannot rule out that aggregates may be permeable at microscales, this shows that the permeability of our field-collected aggregates is too low to affect their hydrodynamics and sinking velocity, despite their high porosities. Yet, even a low degree of permeability, not detectable at the resolution investigated here, could affect solute exchange and microbial colonization. Aggregate impermeability can be explained by high concentrations of transparent exopolymer particles (TEP) that may limit advection, but not diffusion, within aggregates (*16*). TEP is also the most likely reason that large aggregates, albeit being highly porous, are not permeable, as TEP occupy the vast majority of the pore space, and constitute a much larger amount of the “non-fluid” fraction than solid components (*5, 56, 57*). This suggests that natural, *in situ*-formed aggregates are mainly diffusive systems where any mass transport to or from the aggregates has to occur across the diffusive boundary layer.

### Formulating a global scaling relationship between aggregate size and carbon flux

Because of the relative ease of imaging aggregates with bio-optical systems compared to measuring the sinking velocity and carbon content of individual aggregates, accurately estimating carbon flux from images has become an important aspect of marine carbon cycling research. This requires the basic assumption that we can relate both sinking velocity and carbon content to aggregate characteristics, in particular aggregate size. For a long time, it was assumed that aggregate size and sinking velocity scale according to a quadratic power function, but while this generally holds true for laboratory-formed aggregates, it has become increasingly clear that this relationship is much weaker for *in situ* aggregates (*14, 58*). With our approach, we can provide a mechanistic explanation for this observation, because, once we included variable solid hydrated density and our size-to-porosity relationship into the force balance equation, we were able to predict sinking velocity for both laboratory-formed and *in situ*-formed aggregates. The inclusion of porosity and density distinguishes our relationship from those of (*7*) and (*25*), which were shown to over- or underestimate carbon flux depending on region and season, calling into question whether a global scaling relationship between aggregate size and carbon content can be established at all. We argue that they were not globally applicable because of differences in solid hydrated densities between the source communities from which sinking aggregates were formed. If the density of the solid fraction is taken into account, as presented in this study, formulating multiple scaling relationships for different regions and/or seasons should become obsolete.

### Contribution of small aggregates to export fluxes

Once a size-dependent porosity term is included in the force balance equation, it becomes obvious that small aggregates (here defined as aggregates <500 µm, i.e. below the cut-off of what is defined as “marine snow”) contribute even more to carbon export than would be expected without considering size-dependent changes in aggregate porosity. We found that small aggregates, due to their lower porosities and generally higher carbon-to-volume ratios compared to larger aggregates, combined with high aggregate concentrations, on average, contribute 80% of the cumulative carbon flux in our study area, and 67% globally (Fig. 5, D). Studies examining aggregate size-distributions with bio-optical systems have previously established the significance of small aggregates (<500 μm) for carbon export fluxes (*28, 50, 59–61*) and matched our results well (*60*), meaning that we can deliver a mechanistic basis for these observations. Our results further show that the relative contribution of small aggregates to carbon flux is higher in less productive regions, which is in accordance with (*60*), who hypothesize that the apparent correlation with sinking flux can be partially attributed to differences in community structure, such as cell sizes in the initial plankton assemblage. Community size structure is expected to change in most future climate change scenarios (*62*), further highlighting the importance of understanding the respective contribution of aggregates of different sizes to export fluxes. Our results further highlight the importance of camera systems for aggregate profiling over direct measurements, as aggregates <500 μm can only barely be seen with the naked eye. Another advantage of *in situ* camera systems that can be exploited further is that mean grey values of the imaged aggregates can potentially be correlated with their porosities. We extracted the mean grey values (ranging from 0 to 255, with 0 being black and 255 being white) from images of aggregates taken under the microscope and found that the proportion of solid matter increased exponentially with decreasing mean grey value. This conforms to what would intuitively be assumed due to lower amounts of pore space (equaling a low mean grey value) in denser aggregates (Fig. S1, B).

Our measurements provide a quantitative and mechanistic explanation for the previously reported considerable contribution of small aggregates to the biological carbon pump. Small aggregates (<500 μm) contribute on average 67% to carbon export globally, which can be attributed to their high abundance and increased compactness. This does not only increase the size-specific sinking velocity, but also their size-specific carbon content. The contribution of small aggregates to carbon export exhibits regional differences and is highest at mid-latitudes, where the phytoplankton community is usually dominated by diatoms and coccolithophores and compact copepod fecal pellets often contribute substantially to export flux (*63*). These findings have far-reaching ecological implications, such as enhanced vertical connectivity in the ocean, with potential effects on oceanic carbon and nutrient budgets that remain to be explored.

## Materials and Methods

### Experimental Design

The study was carried out in the Northwest African Eastern Boundary Upwelling Ecosystem off the Mauritanian Coast between July 12^th^ and August 19^th^ 2017. Upwelling areas are known for their high primary productivity and export, which is ideal as our objective was to collect a high number of diverse aggregates *in situ*. The sampling was designed in a way that allowed us to maximize the number of parameters measured for each aggregate, including microscopy, PIV, sinking velocity, size, and solid hydrated density. The combination with sediment trap deployments and vertical camera profiles from the same cruise allowed us to relate our findings to *in situ*-processes and infer underlying mechanistic processes driving sinking and export of marine aggregates. Please refer to (*64*) for details about oceanographic parameters measured at the time of the cruise.

### Aggregate collection

Aggregates were collected non-destructively with a Marine Snow Catcher (MSC;(*13*)) at depths between 20-150 m and left to settle on deck for 5-10 h, based on visual observations of when the aggregates sank to the base of the MSC. Hereafter, single aggregates were picked from the base of the MSC with a wide-mouth bore pipette and gently transferred to a vertical flow system to record the size, sinking velocity and flow field around each aggregate. The flow system was modified from (*15*) to optimize laminar flow of the water (Fig. 1, A-C). The water in the vertical flow system was collected from the same MSC deployment as the aggregates, GF/F-filtered (mean pore size ∼0.75 µm), and kept at a constant temperature of 18.2°C (±0.2°C). In addition to collecting natural, *in situ*-formed aggregates in the field, we prepared agar spheres made up of 1% agar (wt/wt) mixed with 1% yeast (wt/wt) dissolved in seawater. They were used as model aggregates for impermeable, perfect spheres where solute and gas exchange between the interior and the ambient water occurs via diffusion only. Spheres were formed by dripping the agar solution into a layer of paraffin oil and letting the spheres settle out into an underlying layer of filtered seawater (*65, 66*). Agar spheres were immersed in water of equal density as used for the experiments for at least 10 minutes before use to allow salt to diffuse between the ambient and pore water until equilibrium was reached.

### Particle Image Velocimetry

For Particle Image Velocimetry (PIV), the water reservoir of the vertical flow system was seeded with polyamide particles with a mean diameter of 5 µm and a density of 1.1 g/cm^3^ (ILA 5150 GmbH) to track the velocity and direction of water flow around the aggregates. The center section of the flow chamber was illuminated with a green LED Pulsing System (LPS, ILA 5150 GmbH) attached to an LED head with optics to create a vertical, 500 µm-wide light sheet illuminating the flow chamber (Fig. 2 D, E). The LPS was pulsed at frequencies ranging from 20-30 Hz depending on the flow velocity. The upward flow in the vertical flow system was adjusted until aggregates remained suspended at a distance of approximately 3-5 aggregate diameters above the net of the vertical flow system to eliminate any interaction between surfaces. A series of at least 150 images was recorded with a camera (µEye CP, IDS Imaging Development Systems GmbH), with an attached macro zoom telecentric lens (computar, TEC-V7X, 47.65 - 106.26 mm). The camera was triggered by the LPS, i.e., the frame rate was automatically adjusted according to the LED pulse frequency.

PIV recordings were analyzed with the software PivView (PivTech GmbH). Pictures were divided into interrogation windows with an area of 32 pixels each and a 50% overlap. It was qualitatively ensured that each interrogation window contained at least five individual PIV beads. Cross-correlation of each interrogation window between two consecutive images was performed using a discrete fast Fourier transform following the Wiener– Khinchin theorem. To maximize spatial resolution, multi-grid refinement analysis was performed (*67*). The result was a 2-dimensional velocity matrix with a size of 16 px × 16 px at a resolution of 7 µm px^-1^. The procedure was repeated for at least 80 consecutive images using a batch script. Further processing was performed in Matlab 2015b (The Mathworks, Inc.). All processing routines are publicly available at https://github.com/SoerenAhmerkamp/snowPIV. Although marine aggregates were mostly stable within the imposed flow conditions, slight movements in the range of a few pixels were observed. To avoid any blurring while averaging, the movement of the aggregate was corrected using a detection algorithm to re-center the aggregate. For this, PIV seeding was filtered from the raw images using a 2-dimensional median filter with a pixel size of 6 px × 6 px. Subsequently, the image was binarized and the edges of the aggregate were traced. The aggregate was cropped and a cross-correlation analysis with the subsequent images was performed to determine the particle movement.

Based on the position of the center of gravity, a velocity matrix stack composed of the 80 instantaneous velocity measurements was created with the centered aggregate. Finally, the velocity matrix stack was averaged and the position of the aggregate was masked. Aggregates that exceeded a movement of 20 pixels in horizontal or vertical direction within 2-3 seconds of every possible image sequence were excluded from further analysis. Fluid boundary layer thickness was determined from the averaged magnitude of the flow field. For that, a horizontal cross-section was extracted along the equator of the aggregate. The undisturbed maximum velocity along the cross-section is equal to the terminal sinking velocity of the aggregate. We then defined the fluid boundary layer as the distance between the aggregate surface and the point where the velocity increases to 90% of the terminal velocity.

### Aggregate properties

We measured properties relating to the sinking, porosity and shape of marine aggregates to relate them to each other and to numerically describe aggregate sinking behavior. Selected properties including sinking velocity, size, shape factors, solid hydrated density and fluid boundary layer thickness were measured directly, while porosity, excess density and drag coefficient were derived as described in the following. Aggregate size was measured in two different ways: i) equivalent circular diameter (ECD) of aggregates was calculated from the two-dimensional projected area of aggregates as determined by the detection algorithm described above, ii) the x-axis diameter was extracted from images of the PIV recording as the diameter of the aggregate that is perpendicular to the falling direction. Because the x-axis diameter has a direct impact on the aggregate flow field and is a common input parameter for many hydrodynamic equations, e.g., the drag coefficient (*40*), we used the x-axis diameter as the length measure for the aggregates throughout this manuscript.

For comparison to previous sinking velocity measurements, sinking velocity of aggregates was also directly determined in the flow chamber by dividing the flow rate by the cross-sectional area of the flow chamber (Fig. 2, E, F). For this, sinking velocity was recorded for aggregates suspended 3-5 diameters above the net at the height where images were taken for PIV, and one diameter above the net as recommended by (*16*) according to (*15*). PIV results indicated that the flow field around the aggregates was affected by the proximity to the surface. Therefore, PIV measurements were recorded with the aggregates suspended at least 3-5 diameters above the net in order to ensure an undisturbed flow field. After recordings in the vertical flow system, aggregates were transferred to Utermöhl chambers (*68*) and imaged using brightfield and phase-contrast microscopy at 100x-400x magnification for high-resolution imaging of aggregate composition, type, and structure. To maximize image resolution, images taken at different depths of field were stacked with Zerene Stacker (Zerene Systems LLC).

The terminal sinking velocity *w* (cm s^-1^) of sinking aggregates in the water column is determined through the balance of forces acting on it. These forces include gravitational, buoyant, and drag forces. The gravitational force is expressed as

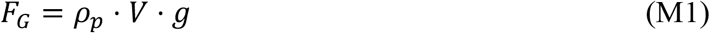

where *𝜌p* is the aggregate density (g cm^-3^), *V* the aggregate volume (cm^3^) and *g* the gravitational acceleration (981 cm s^-1^). At steady state, the gravitational force is counterbalanced by both the buoyancy and drag forces. These forces are respectively given by

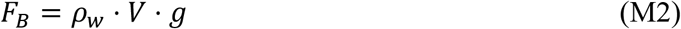

and

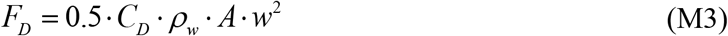

where *ρ*_*w*_ is the water density (g cm^-3^), *A* is the area perpendicular to the settling vector (cm^2^) and *C*_*D*_ is the drag coefficient. For spherical objects sinking at low Reynolds number, (*27*) expressed the drag coefficient as a function of the Reynolds number (*Re*)

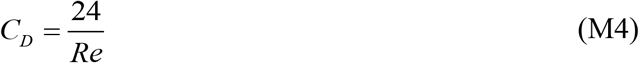

with *Re* defined as

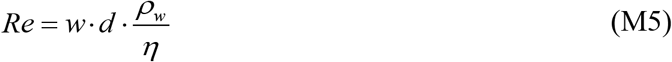

Here *d* and *η* represent the x-axis diameter (cm) and the dynamic viscosity of seawater (g cm^-1^ s^-1^), respectively. By substituting Eqs. M1-M5 into the force balance equation (*F*_*G*_ = *F*_*B*_+*F*_*D*_), the theoretical settling velocity of spherical aggregates is obtained using Stokes’ law (*27*):

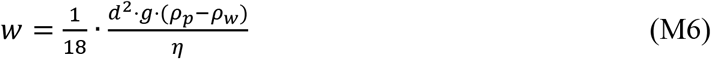

where *𝜌*_*p*_ ™ *𝜌*_*w*_ expresses the aggregate excess density *Δ𝜌*. Using Stokes’ law (Eq. M6), the theoretical sinking velocity *w* is only valid for creeping flow conditions of Re << 1. However, marine aggregates usually fall into an intermediate range of 0.1 < *Re* < 100 (e.g., (*32, 35, 55*)). To increase the range of validity, the drag coefficient can be iteratively determined using the modified drag coefficient (*40*):

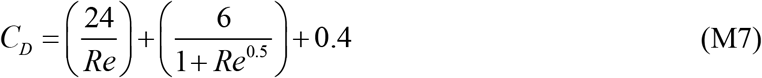

A major challenge associated with predicting marine aggregate sinking velocity is to determine the aggregate excess density. For porous aggregates, the excess density is given as *Δ𝜌* = (*1* ™ *φ) ⋅ 𝜌*_*shd*_ + *𝜌*_*w*_ *⋅ φ* ™ *𝜌*_*w*_, where *φ* is the aggregate porosity, i.e., the ratio of voids to total volume, and *𝜌*_*shd*_ is the solid hydrated density. By applying Eqs. M1-M3 to the force balance equation, the excess density can be expressed as

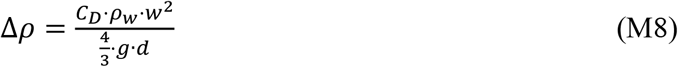

To calculate *Δρ* from Eq. M8, we used the measured sinking velocity and independently determined the drag coefficient. The drag coefficient was derived from the fluid boundary layer thickness *δ*obtained from the PIV flow fields, utilizing a computer simulation-based relationship that correlates the fluid boundary layer to the drag coefficient. This relationship is expressed as:

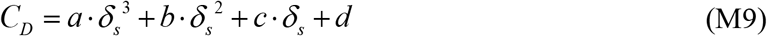

with *a* = 8.8, *b* = −15.1, *c* = 22.3, and *d* = −2.6 and *δ*_*s*_ being the equatorial fluid boundary layer *δ*normalized by the particle x-axis diameter (normal to the flow direction). Eq. M9 was derived through numerical calculations of the drag coefficient from simulated flow fields around spherical aggregates according to (67), and is valid within the range of 0.02 ≤ Re ≤ 20. For each aggregate, we directly measured *ρ*_*shd*_ using the linear gradient method (see section below). Given the measured *ρ*_*shd*_ and calculated *Δ𝜌* (Eq. M8), the porosity of each aggregate was estimated as

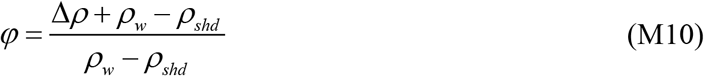

We further characterized the microstructure of the aggregates by determining their 1-& 2-dimensional fractal numbers DF1 and DF2, respectively. Following (*69*), the DF1 and DF2 were calculated from the slopes of log-log relationships of perimeter and area respectively, against the equivalent circular diameter. Particle perimeter, area and diameter were measured from images of the PIV recording using the image processing software FIJI/ImageJ (*70*). To test the impact of aggregate shape on sinking velocities, we also used images from the PIV recording to extract the aspect ratio, area:perimeter, circularity, orientation, convexity, solidity, contour temperature and fractal dimension (collectively termed “shape factors”) with FIJI/ImageJ (*70*) and the ImageJ Shape Filter Plugin (*71*).

The solid hydrated density (*ρ*_*shd*_) of aggregates was determined in a seven-layer density gradient made with dilutions from Ludox TM colloidal silica (Sigma-Aldrich), sucrose and distilled water (following (*45, 72*)) who modified the method from (*73, 74*). The dilutions ranged in density from 1.155 to 1.428 g cm^-3^. Starting with the densest solution, 2 ml of each dilution were gently transferred to 15 ml centrifuge tubes (Corning) and topped with 2 ml of GF/F-filtered seawater. Single aggregates were gently added to the topmost layer of the density gradient using a wide-mouth bore pipette and left to settle into the gradient at 4°C overnight. The following day, 1 ml was sampled from the layer that the aggregate had settled into, which is representative of the density of the solid components of the aggregate. Samples were kept at 4°C in sealed 1 ml-tubes until the end of the cruise, and measured with a density meter (DMA 38, Anton Paar) back in the home laboratory. Because the aggregates were destroyed in the process, this measurement was done at the very end. For sinking velocity and carbon flux predictions, we used the averaged solid hydrated density *ρ*_*shd*_ of each identified aggregate type (phytoplankton aggregates, ballasted mucus aggregates, and degraded fecal pellets).

### Sinking velocity comparisons

To test the suitability of our fitted relationship between aggregate diameter and porosity for predicting sinking velocity, we compared the experimentally measured and the predicted sinking velocities with the porosity included using our best fit between porosity and size, specific for each aggregate type. This combines Eqs. 2.a-2.c and M6 to predict sinking velocity based on the fitted regression between aggregate size and porosity, which was independently derived from the boundary layer-inferred drag coefficient:

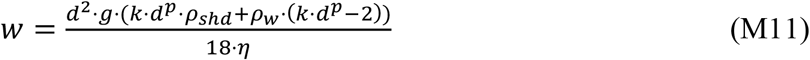

Where *k* is the coefficient and *p* is the exponent of Eq. 2.a. To test the applicability for other datasets from different regions and origins, we collected datasets with known water density and, experimentally measured sinking velocity, size and solid hydrated density *ρ*_*shd*_ and used our relationship to calculate porosity, based on which excess density and sinking velocity were consequently calculated (Eq. M11).

### Sediment trap deployment and aggregate-specific carbon content

To measure *in situ* carbon fluxes, we deployed two drifting sediment traps for a duration of 22 h each. The drifting sediment traps consisted of one trap station each at depths of 100 and 200 m equipped with four collection cylinders. Material from one trap tube per depth was fixed with 1.4% mercury chloride and stored at 4°C for later determination of mass fluxes of carbon, nitrogen, biogenic opal, calcium carbonate, and lithogenic (non-biogenic) material. In trap 1, two collection cylinders per depth were equipped with a viscous cryo-gel (Tissue-Tek, Sakura Finetek) to structurally preserve aggregates in order to determine their abundance and size spectra, and in combination with measured carbon fluxes, to determine size-specific carbon content. Aggregates collected in the gel cups were imaged on board, and images were subsequently processed with the image processing software FIJI/ImageJ (*70*).

Aggregate sizes and numbers across the water column were recorded with the Underwater Vision Profiler 5hd (UVP 5hd; HYDROPTIC, France; (*22*)) in a total of 72 profiles. The UVP was mounted inside the CTD frame and operated in autonomous pressure mode. Images were acquired with a maximum frequency of 20 Hz during every downcast. The sampling volume was approximately 1 L, with a particle quantification threshold of 2 pixels (∼80 µm ESD). Post-processing was accomplished using the ImageJ based software Zooprocess (*75*), as well as the Ecotaxa platform and the Ecopart export tool (*76*). We re-interpolated the data onto 1 m steps within the depth range of 20 m to 500 m. Subsequently, we categorized the aggregates into the same size bins as above. We then calculated the average of all N=72 profiles and integrated the size-specific carbon fluxes. This integration allowed us to calculate depth profiles of carbon fluxes

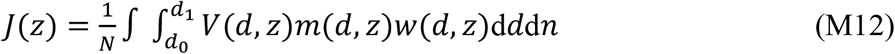

Where *z* is water depth (m), *w(d*) the aggregate settling velocity, *d*_*0*_ = 128 µm and d_*1*_ = 2024 µm are the lower and upper limits for integration, and *m(d,z*) is the particle specific carbon content. The settling velocity and carbon content was calculated by applying our newly derived size-porosity relationship (Eq. 2.a), combined with Eq. M6 (giving Eq. M11).

### Global contribution of small aggregates to export flux

We obtained a comprehensive dataset of 13,291 global under-water-vision (UVP)-based aggregate profiles from the publicly available ECOTAXA database (Last visit: 7/21/2023). After an initial quality check and filtering for completeness of parameters (temperature, salinity) and profile depth, we were left with 10,907 high-quality underwater aggregate profiles from which we extracted the diameter and number of aggregates. We used Eqs. 4, M6 and M12 to calculate the carbon flux at the base of the annual mean mixed layer (*49*). The seawater density and kinematic viscosity were calculated based on temperature and salinity based on empirical relationships (*77*), the porosity was calculated based on Eqs. 2.a-2.c, and the solid hydrated densities were based on literature values for different ocean regions (Table S1). For our measurements, we found a remarkably strong direct relationship between porosity and excess density: *Δρ* = 0.08 · (1 - *φ*)^0.73^, which we employed for the Atlantic Ocean. To gain insight into the impact of small aggregates, we calculated the relative contribution of aggregates ranging from 128 µm to 500 µm to the total carbon flux.

## Acknowledgments

We thank the captain and crew of the B/O Sarmiento de Gamboa and the technical staff of CSIC Unidad de Tecnología Marina (UTM) for their invaluable support at sea, and Anna Belcher for her contribution to the DY050 dataset. CMF further thanks Prof. Adi Torfstein for supporting the finalization of the manuscript.

## Funding

Spanish National Plan for R&D project FLUXES (CTM2015-69392-C3-1-R), co-financed with FEDER funds (JA, XAS)

Helmholtz Program-Orientated Funding (POF IV) topic 6 (Marine Life) and sub-topic 6.3 (The Future Biological Carbon Pump) (MHI, RK)

HGF Young Investigator Group SeaPump “Seasonal and regional food web interactions with the biological pump”: VH-NG-1000 (CMF, MHI)

AWI Strategy Fund Project ‘Climate change impact on ecosystem structure and the efficiency of the biological carbon pump for Antarctic diatom- and *Phaeocystis*-dominated phytoplankton communities (EcoPump)’ (CMF, MHI)

DFG-Research Center/Cluster of Excellence “The Ocean Floor - Earth’s Uncharted Interface”: EXC-2077-390741603 (MHI)

DFG project GR1540/28-1 (HPG)

DFG project GR1540/37-1 (HPG)

DFG project IV124/3-1 (HPG)

MARMA project by the Max Planck Society (SA, JM)

MATIC project by Novo Nordisk Fonden (SA)

Make Our Planet Great Again grant from the French National Research Agency (ANR) within the Programme d’Investissements d’Avenir #ANR-19-MPGA-0012 (RK)

Heisenberg Programme of the German Science Foundation #KI 1387/5-1 (RK)

## Author contributions

Conceptualization: CMF, SA, MHI

Method development and testing: CMF, SA, CH, IK, NM, MHI

Fieldwork: CMF, IK, XAS, JA

Data processing: CMF, SA, RK

Visualization: CMF, SA

Consultation: NM, JM, RK, MHI

Supervision: MHI, AK, HPG, SA

Writing—original draft: CMF, SA, MHI

Writing—review & editing: CMF, SA, MHI, RK, XAS, JM, IK …

## Data and materials availability

All data are available in the main text or the supplementary materials. Complementary data and images are publicly available in the Pangaea data repository. All PIV processing routines are publicly available at https://github.com/SoerenAhmerkamp/snowPIV.

