## Supplementary Material for "Rethinking sinking: Imaging flow fields of natural marine aggregates to infer porosity-dependent changes in sinking velocity and carbon flux"

**The PDF file includes:**

Figs. S1 to S5
Tables S1 to S4

**Other Supplementary Materials for this manuscript include the following:**

Data S1 to S2

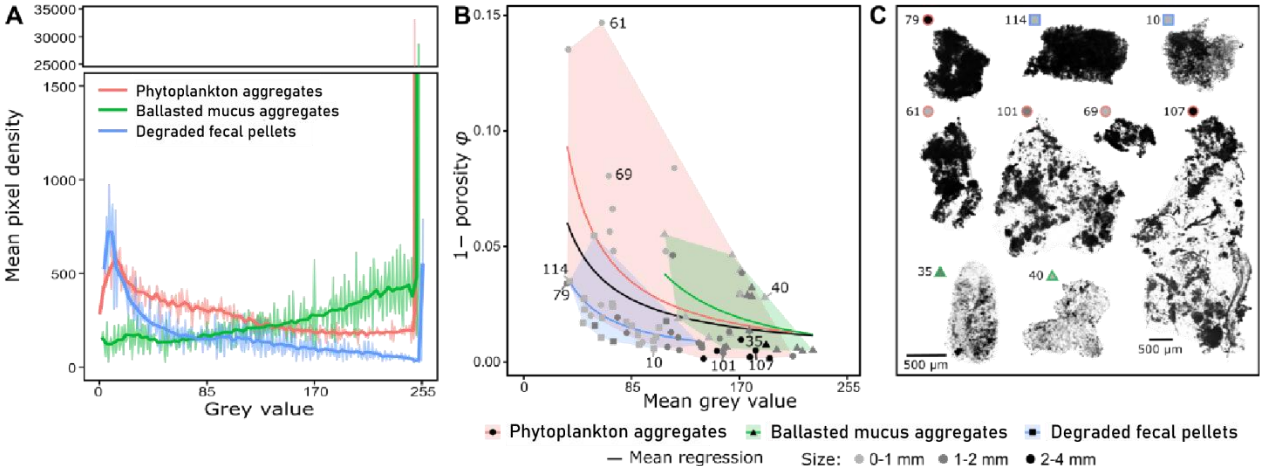


**Fig. S1. Analysis of grayscale microscope images. (A)** Gray value distribution showing distinct patterns for each aggregate type, allowing categorization based on grayscale images alone, **(B)** Correlation between fraction of solid matter and mean grey value for the different aggregate types, including examples **(C)**. The scale bar (bottom left) in panel **C** applies to all aggregates except for aggregate 107, which has a separate scale bar.

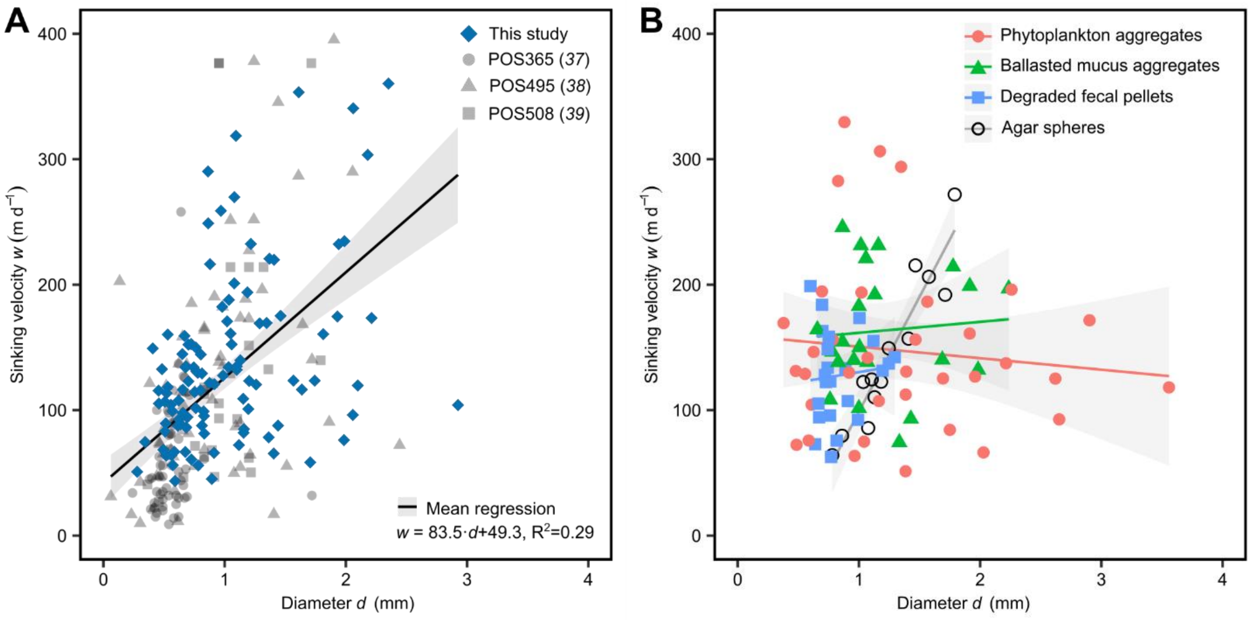


**Fig. S2. Sinking velocity comparisons. (A)** Size-to-sinking relationship of aggregates collected for this study (deep blue) compared to other aggregates collected in the Mauritanian upwelling area (gray) (*37*–*39*). **(B)** Size-specific sinking velocities of model agar spheres ($R^2$=0.89) and aggregates ($R^2$=0.01) collected for this study, categorized by aggregate type.

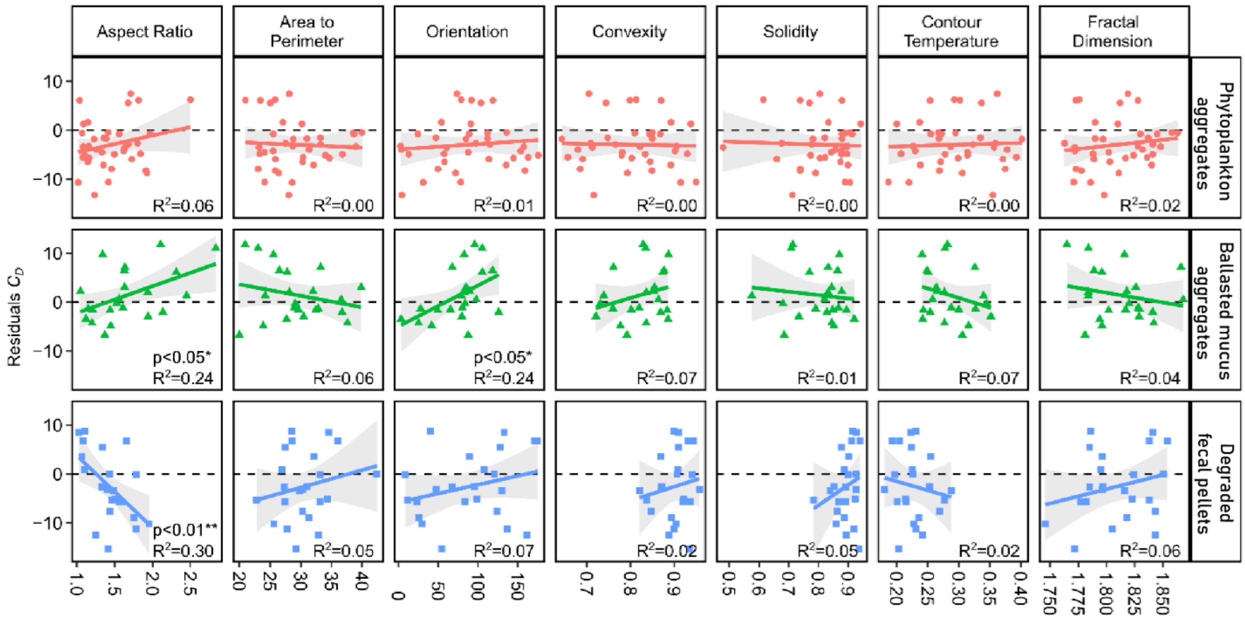


**Fig. S3. Shape analysis.** Drag coefficient residuals for different shape factors categorized by aggregate type.

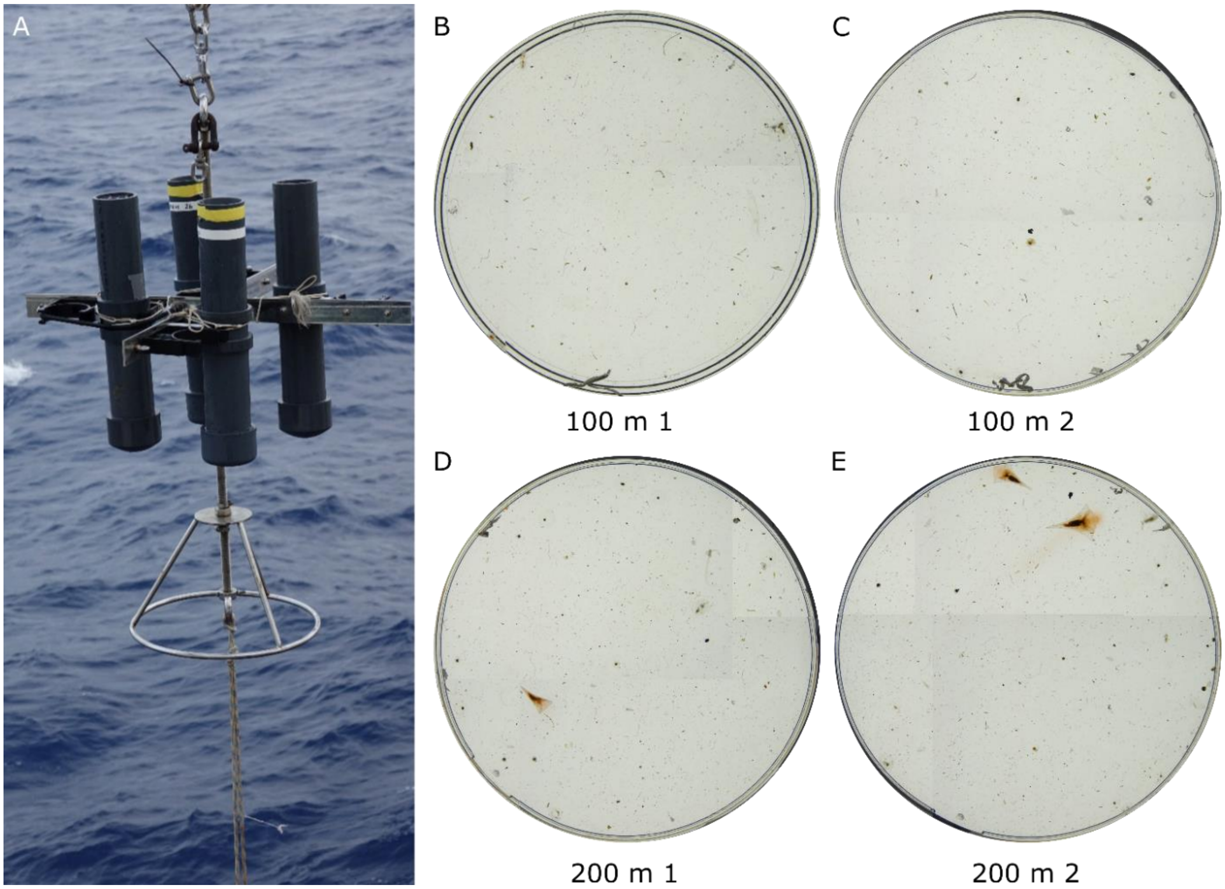


**Fig. S4. Gel trap images**. **(A)** Sediment trap deployment at Station 4 with two sets of gel traps deployed at 100 m **(B, C)** and 200 m **(D, E)**, respectively.

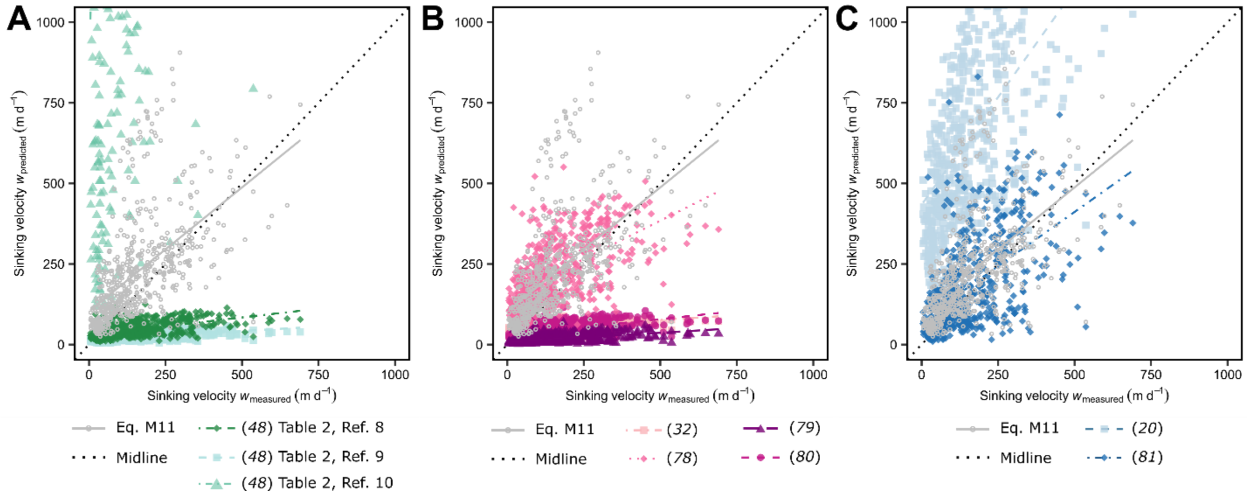


**Fig. S5. Predicted vs. measured sinking velocities using our relationship (grey) and relationships compiled in Table 2 of** (*48*)**.** Comparison to **(A)** empirical relationships established in (*48*), **(B)** *in situ*-formed aggregates (*32*, *78–80*), and **(C)** roller tank-formed aggregates (*20*, *81*). Y-axes in panels **(A)** and **(C)** were cut off at 1000 m, for better visualization.

**Table S1.** Overview of solid hydrated densities of selected aggregate types. (*) marks a sample size of n=1.

| Particle Type | Content/origin | Solid hydrated density (g cm$^{-3}$) | Origin | Reference |
|---|---|---|---|---|
| **Diatom aggregates** | unspecified | 1.17±0.03 | NW Africa | *This study*; Data S2 |
| | unspecified | 1.08 | Southern California Bight | (*12*) |
| | *S. marinoi* | 1.18±0.12 | Culture | (*16*) |
| | *S. marinoi* | 1.15±0.02 | Culture | (*43*) |
| | *P. antarctica, C. debilis, P. turgidula* | 1.15±0.01 | Culture | (*44*); Data S2 |
| | *Chaetoceros spp., Skeletonema spp., Thalassiosira spp., Coscinodiscus spp.* | 1.26±0.10 | North Sea | (*45*) |
| | *S. marinoi, T. pseudonana. T. weissflogii* | 1.09±0.03 | Culture | (*46*); Data S2 |
| | Marine snow | 1.17±0.15 | Porcupine Abyssal Plain | (*47*); Data S2 |
| | *T. weissflogii* | 1.17±0.02 | Culture | (*72*) |
| | *Chaetoceros* sp. | 1.31±0.03 | Culture | (*82*) |
| | Marine snow | 1.34±0.11 | Bedford basin | (*82*) |
| | *Phaeocystis spp., Pseudo-nietzschia spp., Chaetoceros spp., Fragilariopsis spp., Corethron spp., Coscinodiscus spp.* | 1.12±0.02 | Weddell Sea | (*83*); Data S2 |
| | *Synedra* sp. | 1.284±0.00 | Culture | (*84*) |
| **Cyanobacteria aggregates*** | unknown | 1.075 | NW Africa | *This study*; Data S2 |
| **Coccolithophore aggregates** | *E. huxleyi* | 1.11 | Culture | (*16*) |
| | *E. huxleyi* | 1.14±0.05 | Culture | (*55*) |

| | | | | |
|---|---|---|---|---|
| | *E. huxleyi* | 1.19±0.03 | Culture | (*85*) |
| **Copepod fecal pellets** | *T. longicornis* | 1.60 | Culture | (*16*) |
| | *A. tonsa* | 1.15±0.03 | Culture | (*74*) |
| | *C. helgolandicus* | 1.26±0.01 | Culture | (*86*) |
| | Small copepods | 1.28±0.11 | Monaco coast | (*87*) |
| | *A. patersoni* | 1.15±0.05 | Monaco coast | (*87*) |
| **Euphausiid fecal pellets** | *M. norvegica* | 1.22 | Bahamas | (*88*) |
| | *M. norvegica, E. krohnii, N. megalops* | 1.29±0.1 | Ligurian Sea | (*89–91*) |
| | unspecified | 1.23 | Monaco coast | (*91*) |
| **Salp fecal pellets** | *S. fusiformis, P. socialis* | 1.10 | California Current | (*92*) |
| | unspecified | 1.275±0.05 | Santa Barbara Basin | (*93*) |
| | unspecified | 1.19±0.03 | Unknown | (*94*) |
| **Appendicularian fecal pellets** | unspecified | 1.70 | NW Africa | (*16*) |
| **Degraded fecal pellets** | unknown | 1.31±0.08 | NW Africa | *This study*; Data S2 |
| **Ballasted mucus** | unknown | 1.26±0.05 | NW Africa | *This study*; Data S2 |
| **TEP** | | 0.70-0.84 | Santa Barbara Channel | (*95*) |

**Table S2.** Range of maximum aggregate length, number of samples, fractal dimensions (DF1 and DF2, respectively, ± standard deviation) and associated $R^2$ for all aggregate types.

| Particle type | Size range | n | DF1 | $R^2$ | DF2 | $R^2$ |
|---|---|---|---|---|---|---|
| All field-collected aggregates | 0.4-3.5 mm | 79 | 1.43(±0.22) | 0.89 | 1.79(±0.20) | 0.93 |
| Phytoplankton aggregates | 0.6-3.5 mm | 36 | 1.47(±0.24) | 0.91 | 1.91(±0.23) | 0.94 |
| Ballasted mucus aggregates | 0.4-2.0 mm | 20 | 1.11(±0.22) | 0.82 | 1.62(±0.21) | 0.90 |
| Degraded fecal pellets | 0.7-2.3 mm | 23 | 1.47(±0.16) | 0.88 | 1.81(±0.16) | 0.91 |
| Agar spheres | 0.8-2.0 mm | 13 | 1.18(±0.15) | 0.82 | 2.00(±0.05) | 0.99 |

**Table S3.** Details of sediment drifting traps deployed during the Fluxes I cruise (*na*=not available).

| | Trap 1 | Trap 2 |
|---|---|---|
| Station | 4 | 12 |
| Date (dd/mm/yyyy) | 16/07/2017 | 20/07/2017 |
| Coordinates (lat, lon) | 23.0, -19.3 | 23.0, -26.0 |
| Deployment time (h) | 22 | 22 |
| Flux $POC_{100\ m}$ (mg $m^{-2}$ $d^{-1}$) | 42 | 63 |
| Flux $POC_{200\ m}$ (mg $m^{-2}$ $d^{-1}$) | 42 | 51 |
| $C{:}N_{100\ m}$ | 10 | 5.9 |
| $C{:}N_{200\ m}$ | 10.2 | 6.7 |
| Gel traps deployed? | Yes | No |
| Flux $Abundance_{100\ m}$ ($m^{-2}$ $d^{-1}$) | $5.53\times10^{7}$ | *na* |
| Flux $Abundance_{200\ m}$ ($m^{-2}$ $d^{-1}$) | $7.61\times10^{7}$ | *na* |
| Flux $Volume_{100\ m}$ ($mm^3$ $m^{-2}$ $d^{-1}$) | $1.96\times10^{7} \pm 7.49\times10^{5}$ | *na* |
| Flux $Volume_{\ 200\ m}$ ($mm^3$ $m^{-2}$ $d^{-1}$) | $2.29\times10^{7} \pm 7.28\times10^{5}$ | *na* |
| Aggregate $size_{100\ m}$ (µm) | 548 ± 428 | *na* |
| Aggregate $size_{200\ m}$ (µm) | 547 ± 389 | *na* |

**Table S4. Comparison between measured and predicted sinking velocities.** Size *d* and sinking velocity *w* (mean±standard deviation) of aggregates for which *w* was predicted using our size-dependent porosity ($\varphi$) relationship.

| **Reference** | | ***n*** | ***d* (mm)** | $w_{measured}$ **(m d$^{-1}$)** | $w_{predicted}$ **(m d$^{-1}$)** | $w_{predicted}$ **/**$w_{measured}$ |
|---|---|---|---|---|---|---|
| This study | Phytoplankton aggregates | 36 | 1.4±0.8 | 147.2±68 | 206.6±67 | 1.15 |
| | Ballasted mucus aggregates | 22 | 1.2±0.4 | 221.5±5 | 68.2±26 | 1.25 |
| | Degraded fecal pellets | 23 | 0.8±0.2 | 128.3±36 | 126.2±18 | 0.91 |
| (*43*) | | 176 | 3.5±1.3 | 231.2±136 | 258.2±115 | 1.49 |
| (*44*) | | 39 | 4.1±1.9 | 174.8±119 | 180.7±58 | 0.84 |
| (*45*) | | 160 | 2.0±0.9 | 137.4±90 | 491.0±248 | 1.97 |
| (*46*) | | 92 | 1.5±0.8 | 115.3±69 | 121.2±39 | 0.85 |
| DY050 (*47*) | | 109 | 0.7±0.5 | 66.6±80 | 68.2±26 | 0.43 |

**Data S1. Unpublished data for Figs. 5, A; S2, A.** Equivalent spherical diameter $d_{ESD}$ and sinking velocity $w$ of roller tank-formed and *in situ*-formed aggregates from unpublished datasets.

**Data S2. Unpublished data for Table S1.** Solid hydrated density $\rho_{shd}$ of aggregates from unpublished datasets presented in Table S1.
